# Multi-scale predictive modeling discovers Ibudilast as a polypharmacological agent to improve hippocampal-dependent spatial learning and memory and mitigate plaque and tangle pathology in a transgenic rat model of Alzheimer’s disease

**DOI:** 10.1101/2021.04.06.438662

**Authors:** Giovanni Oliveros, Charles H. Wallace, Osama Chaudry, Qiao Liu, Yue Qiu, Lei Xie, Patricia Rockwell, Maria Figueiredo-Pereira, Peter A. Serrano

**Affiliations:** Ph.D. Program in Biochemistry, The Graduate Center, CUNY, 10016; Department of Biological Sciences, Hunter College, CUNY, 10065; Department of Computer Science, Hunter College, CUNY, 10065; Ph.D. Program in Biology, The Graduate Center, CUNY, 10016; Ph.D. Program in Computer Science and Biochemistry, The Graduate Center, CUNY, 10016; Department of Psychology, Hunter College, CUNY, 10065; Helen and Robert Appel Alzheimer’s Disease Research Institute, Feil Family Brain & Mind Research Institute, Weill Cornell Medicine, Cornell University, 10021

## Abstract

Alzheimer’s disease (AD) is a multifactorial disease that exhibits cognitive deficits, neuronal loss, amyloid plaques, neurofibrillary tangles and neuroinflammation in the brain. We developed a multi-scale predictive modeling strategy that integrates machine learning with biophysics and systems pharmacology to model drug actions from molecular interactions to phenotypic responses. We predicted that ibudilast (IBU), a phosphodiesterase inhibitor and toll-like receptor 4 (TLR4) antagonist, inhibited multiple kinases (e.g., IRAK1 and GSG2) as off-targets, modulated multiple AD-associated pathways, and reversed AD molecular phenotypes. We address for the first time the efficacy of ibudilast (IBU) in a transgenic rat model of AD. IBU-treated transgenic rats showed improved cognition and reduced hallmarks of AD pathology. RNA sequencing analyses in the hippocampus showed that IBU affected the expression of pro-inflammatory genes in the TLR signaling pathway. Our results identify IBU as a potential therapeutic to be repurposed for reducing neuroinflammation in AD by targeting TLR signaling.

## Introduction

Alzheimer’s disease (AD) currently affects about 5.8 million Americans, the majority of which are over the age of 65, with a subpopulation of patients under this age cutoff. Experts speculate that by 2050, expenditure set aside for AD will exceed $2.8 trillion and over 14 million people over the age of 65 will be affected^1^. The hallmarks of this disease consist of neuronal loss, amyloid plaques, neurofibrillary tangles, and neuroinflammation detected in the brain of AD patients^2^. In addition, cognitive deficits associated with AD include loss of memory and progressive impairment of thought and reasoning. However, the catalysts for setting AD into motion are still being investigated. Aside from the aforementioned changes, evidence has emerged to suggest that there is a genetic predisposition to AD that leaves vulnerable populations more at risk, with environmental factors and lifestyle possibly playing a role in disease progression^3^.

FDA-approved drugs designed to treat AD through targeting plaques or tangles, do not halt disease progression, having a reported 99.6% failure rate^4^. This failure can be attributed to the complexity of AD, not just the plaques or tangles^2^. One such factor contributing to the pathology of AD is neuroinflammation. This occurs when there is cellular damage in the brain, leading to recruitment of glial cells (microglia and astrocytes), to clear the damage and restore neuronal function^5^. However, chronic neuroinflammation resulting from persistent activation of pro-inflammatory responses in combination with delays in anti-inflammatory responses, causes buildup of cellular debris and neuronal damage. Selecting neuroinflammation as a therapeutic strategy to combat AD may prove beneficial, considering the promises it has shown with treating other neurodegenerative disorders, such as amyotrophic lateral sclerosis (ALS) and multiple sclerosis (MS)^6, 7^. Therapeutics aimed at these two disorders showed promise by amplifying the neuroprotective properties of glial cells, a strategy that could be mimicked by targeting AD pathology from the neuroinflammatory perspective. There is an urgent need to study mechanisms currently in place to combat other neurodegenerative diseases and adapt their strategies to target AD. The major benefit to this approach would be the likelihood that disease progression could be slowed or even halted.

We focused on modulating neuroinflammation by targeting the toll-like receptor (TLR) signaling pathway to combat AD pathology. Toll-like receptors (TLRs) are a family of proteins on the surface of cells that recognize molecular structures that are largely shared by products released by pathogens, and then initiate an appropriat response such as inflammation^8^. TLR signaling pathways are stimulated by pathogen-associated molecular pattern molecules (PAMPs), such as lipopolysaccharides (LPS) produced by Gram-negative bacteria^9^. Damage-associated molecular pattern molecules (DAMPs), like amyloid-beta peptide (A*β*) can also stimulate the pathway. Upon TLR stimulation, myeloid differentiation primary response protein 88 (MyD88) is recruited to the receptor along with interleukin-1 receptor associated kinases (IRAKs)^10^. A fully-assembled myddosome complex is then formed, resulting in IRAK1 activation through autophosphorylation and subsequent dissociation from the myddosome^11^. In turn, IRAK1 activates tumor necrosis factor (TNF)-receptor associated factor 6 (TRAF6), a ubiquitin ligase. TRAF6 ultimately promotes activation of the transcription factor nuclear factor kappa B (NF*κ*B)^12^. NF*κ*B then stimulates transcription of genes, such as TNF*α*, IL1, IL6 and IL18, encoding for pro-inflammatory cytokines^13^. The TLR pathway appears to be an ideal target for therapeutic intervention, as it is central to the pro-inflammatory cascade involved in neuroinflammation.

To identify existing drugs that can modulate neuroinflammation in AD, we applied a predictive modeling framework that integrates machine learning with biophysics and systems pharmacology to repurpose approved drugs for AD treatment with the focus on anti-inflammatory agents. We predicted that Ibudilast (IBU), a nonselective phosphodiesterase inhibitor and a TLR4 antagonist^14 15^, inhibits IRAK1 as an off-target, modulates multiple AD-associated pathways, and reverses AD patient’s molecular phenotype. IBU is a Japanese medication previously recommended for the treatment of post-stroke complications and asthma^16^. The primary mode of action of IBU is by way of neuroprotection and anti-inflammatory effects. In a clinical trial for opioid withdrawal, IBU was shown to be a TLR4 antagonist and to reduce glial cell activation^17^. Moreover, IBU has shown promise in clinical trials against multiple sclerosis (MS) and amyotrophic lateral sclerosis (ALS)^18^.

As a phosphodiesterase inhibitor, IBU is able to maintain cyclic AMP levels and inhibit T-cell immune function^19^. Furthermore, IBU helps promote IL10 production, while simultaneously inhibiting the production of TNF*α* and NO^20^. A phase two clinical trial for progressive MS tested oral administration of IBU (≤ 100 mg daily) or placebo for 96 weeks^21^. After treatment, most patients exhibited less brain atrophy than the placebo-treated patients. These results were significant, provided that intervention occurred in the early stages of the disease^22^. Based on this study, we evaluated the effects of long-term (six-months) treatment of IBU in a transgenic rat model of AD (Tg-AD) for improvement of spatial memory performance and mitigation of hippocampal AD pathology.

## Results

### IBU is predicted to be a potential drug candidate for AD treatment

Given that AD is a multi-genic systematic disease and there are no validated drug targets and effective therapeutics, conventional one-drug-one-target drug discovery process could be less fruitful. Thus, we sought to discover polypharmacological agents that can modulate multiple pathological processes of AD.

We applied a multi-scale predictive modeling approach to repurposing existing anti-inflammatory drugs for potential AD treatment. The premise of our approach is based on a systematic view of drug actions. Drugs commonly interact with not only their intended protein target (i.e., on-target), but also multiple of other proteins (i.e., off-target). These on-targets and off-targets collectively induce phenotypic response of biological systems via biological networks, which can be characterized by transcriptomics. Thus, a successful compound screening requires the deconvolution of drug actions on a multi-scale, from molecular interactions to network perturbations.

We ranked drugs by their ability to reverse the gene expression profile of AD patients^23^. We first obtained the differential gene expression profile of microglia from a group of AD patients vs healthy controls in the AMP-AD data portal^24^, and considered it as the molecular phenotypic signature of AD (supplemental material Table S1). Then we compared the drug-induced gene expression profile of approximately 20,000 drugs (dosage < 1 μM) with the AD signature such that the downregulated and upregulated genes in AD will be upregulated and downregulated by the drug, respectively. On the top 100 ranked drugs (supplemental material Table S2), three of them are anti-inflammatory drugs, as shown in Table 1. Because sulfasalazine and sasapyrine may not be able to permeant blood-brain-barrier (BBB), we focused our studies on IBU.

**Table 1.**
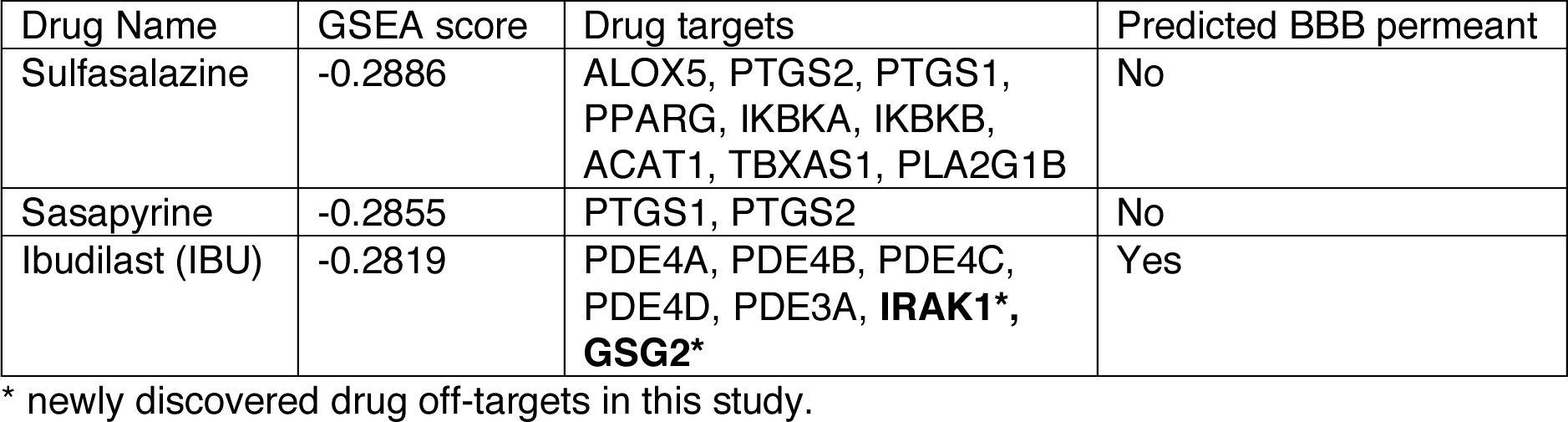
Top 3 ranked anti-inflammatory drugs that are predicted to reverse the gene expression profile of certain AD patients

Using a structure-augmented machine learning method for predicting genome-scale drug-target interactions, we predicted that protein kinases were the off-target of PDE3A inhibitors^25^. The KinomeScan^TM^ assay confirmed our predictions. IRAK1 and HASPIN (GSG2) are effectively inhibited by IBU under the concentration of 10 μM with a percentage control of 6.8 and 9.4, respectively (Supplemental material Table S3). IRAK1 is involved in the TLR and interleukin-1 signaling pathways, and plays a key role in regulating inflammation. Genome-Wide Association Studies (GWAS) discovered that GSG2 is located in the locus associated with AD^26^. Under a high concentration of 100 μM, IBU inhibits additional multiple kinases that are associated with AD and inflammation, such as PIK3C3, CDK4, JNK1, JNK3, HIPK2, HIPK3, TAOK1, TAOK3 (Supplemental material Table S4).

The hypothesis that IBU modulates multiple AD-associated pathways was further supported by text mining. We first recognized all terms that were chemicals, genes or pathways, and diseases in the PubMed abstracts. Then we applied word2vec^27^, a machine learning technique, to represent all terms as vectors that encode their semantic relationships^28^. Finally, the relationships between terms were quantified by the cosine similarity between term vectors.

Using this method, we found that IBU was associated with multiple AD pathological processes including TLR/MYD88/NF*κ*B pathways, TNF pathway, lipopolysaccharide synthesis pathway, and herpes simplex virus infection with the false discovery rate (FDR) less than 1.0e-3.

Altogether, IBU could be an excellent lead compound to design polypharmacological drugs for the treatment of AD by targeting multiple pathological processes.

### IBU improves spatial learning and memory

The hippocampal-dependent, active place avoidance (aPAT) behavioral task was performed at 11 months of age on Tg-AD rats and their wild type (WT) littermates (Fig 1A). Male and female rat performances were combined and four groups were analyzed: wild-type untreated (WTNT), wild type treated with IBU (WTTR), transgenic untreated (TGNT), and transgenic treated with IBU (TGTR). For each trial, data were analyzed using two-way ANOVA with Sidak’s post-hoc test across all six training trials for latency to first entrance into shock zone. To determine whether short-term working memory deficits were more pronounced during the acquisition phase (trials 1-3) vs asymptotic performance phase (trials 4-6) of training, we analyzed these phases separately. Our results show that at 11 months of age WTNT performed significantly better during acquisition (trials 1-3) compared to TGNT (F_(1,26)_ = 3.578, p = 0.0359) with post-hoc differences at trials 2 and 3 (t = 3.051, p = 0.0402 and t = 2.842, p = 0.0498 respectively; Fig 1B). Following 6 months of IBU treated chow or control chow, TGTR show significant improvement during acquisition compared to TGNT controls (F_(1,27)_ = 4.603, p = 0.0411) with post-hoc differences at trial 3 (t = 2.898, p = 0.0486; Fig 1D). This enhanced performance was not observed between WTNT and WTTR conditions (F_(1,28)_ = 2.525, p = 0.1233), suggesting beneficial effects of IBU-treatment under pathological conditions (Fig 1E). TGTR rats performed equivalently to WTTR controls (F_(1,29)_ = 0.4481, p = 0.5085; Fig 1C). There were no significant effects observed during asymptotic performance. Fig 1F shows the tracking for individual rats during trial 3 across the four treatment conditions. 24h after the last training trials rats were tested for avoidance of the shock zone in the absence of shock. The results show that IBU treated rats performed better than controls for max latency to avoid the shock zone, regardless of genotype (F_(1,55)_ = 4.033, p = 0.0495) (Fig 1G).

**Figure 1.**
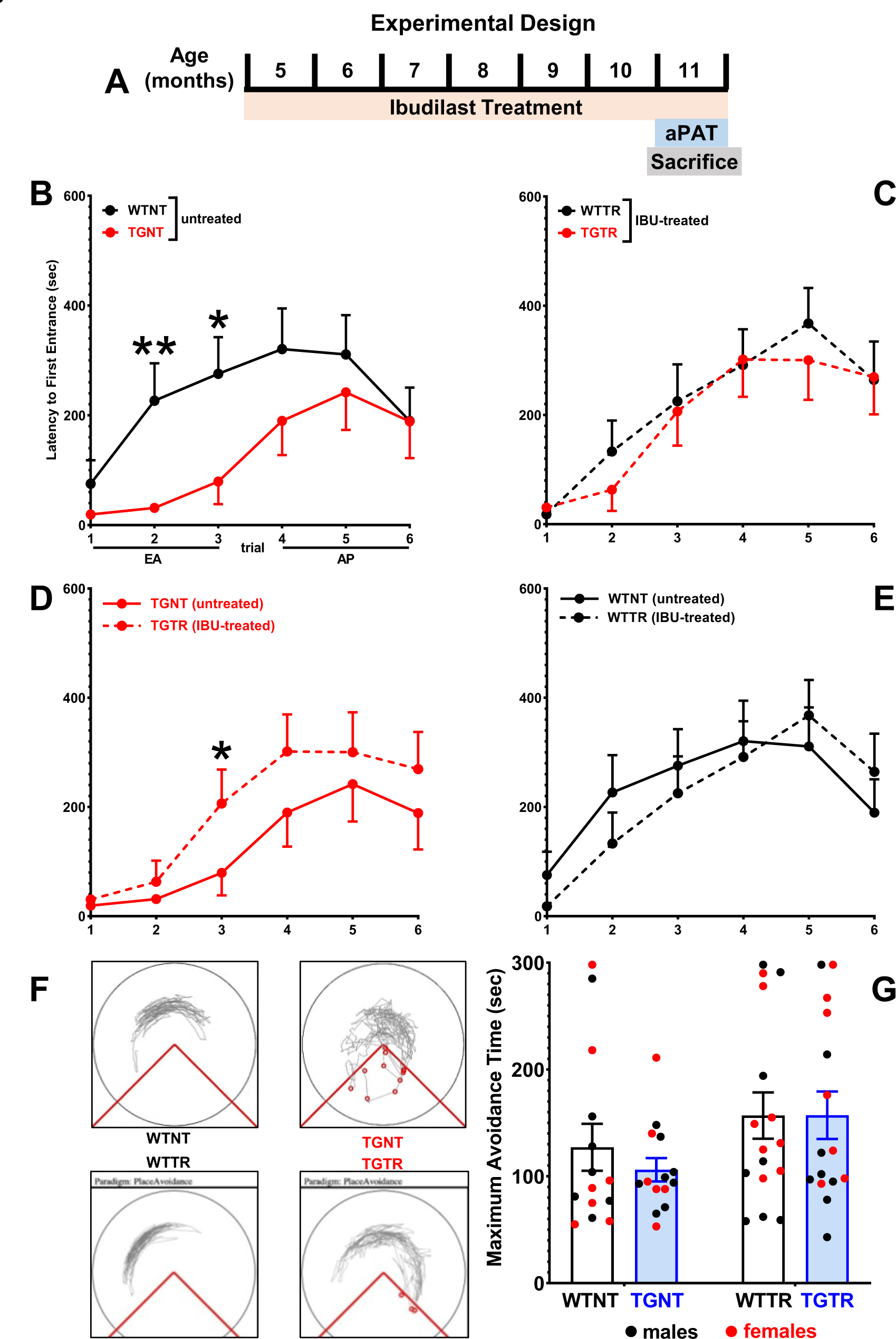
Cognitive effects of IBU-treatment on spatial learning and memory performance. Experimental timeline (A). Latency to first entrance during training (B-E). Early acquisition (EA) shows significant differences between WTNT vs TGNT (B). TGTR perform significantly better than TGNT during EA (D). No differences in EA between WTNT vs WTTR (E) and WTTR and TGTR (C). No differences were observed in asymptotic performance (Trials 4-6) across all comparisons (B-E). Track tracing of individual rat performance for trial 3 across treatment conditions (F). Test data show overall significant improvement in maximum time to avoid during test following IBU treated vs untreated (G). *p < 0.05, **p < 0.01.

### IBU significantly reduces Aβ plaque burden in the dentate gyrus

We evaluated the presence of A*β* plaques in the hippocampus and within discrete subregions (Fig 2A-B), between IBU treated and untreated Tg-AD rats. Our results show that A*β* plaques following IBU-treatment were not significantly different from untreated Tg-AD rats in the hippocampus collapsed across subregions (Fig 2C; t = 0.08131, p = 0.9359). Analyses of different subregions separately shows no significant effects in CA1 (Fig 2D; t = 0.5707, p = 0.5735), CA3 (Fig 2E; t = 0.3351, p = 0.7405) and subiculum (SB) (Fig 2G; t = 0.3020, p = 0.7652). Significant reduction in A*β* plaque load was identified in the dentate gyrus (Fig 2F; t = 3.449, p = 0.0021).

**Figure 2.**
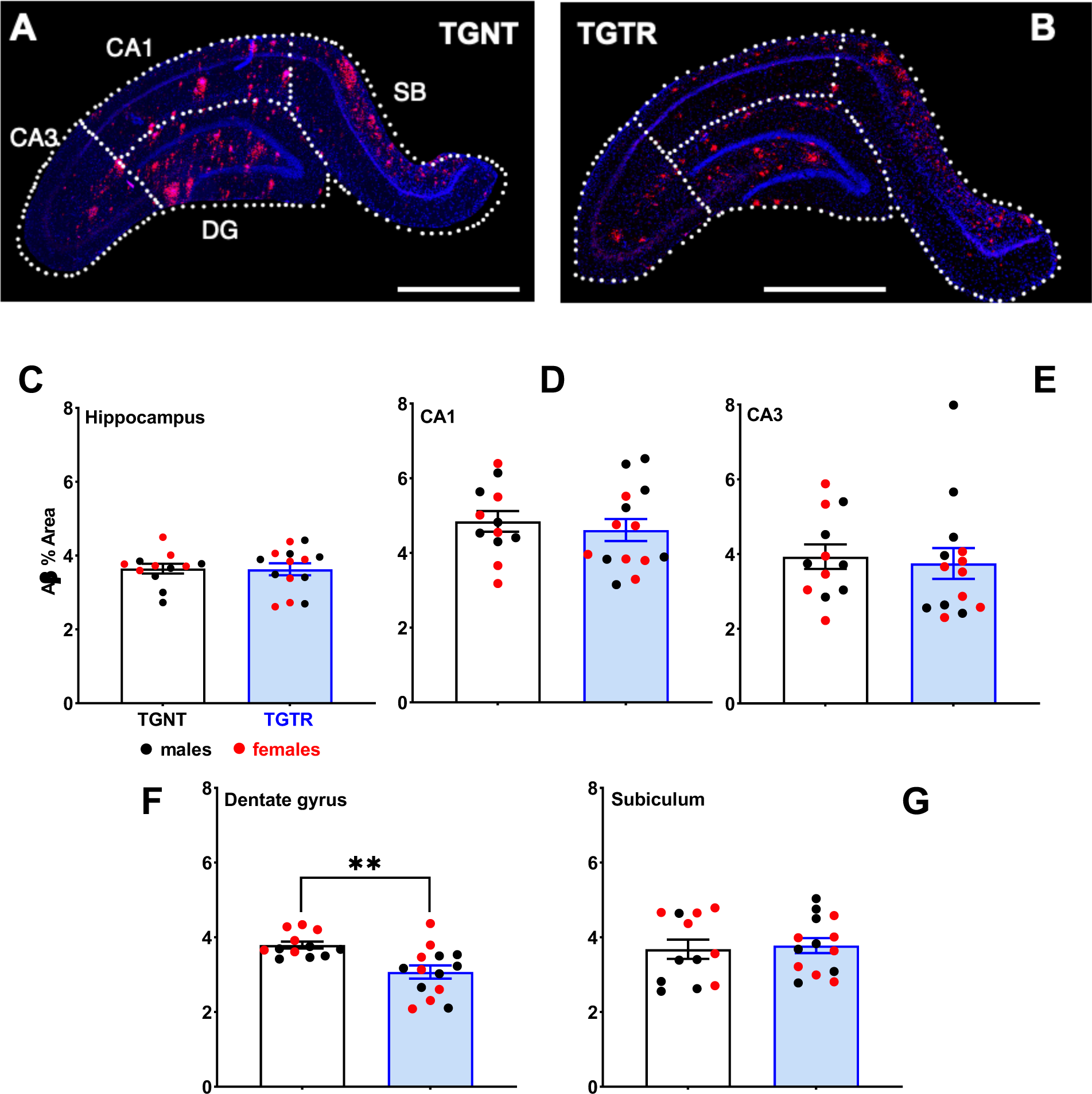
A*β* plaque load is differentially reduced across hippocampal subregions. IHC for A*β* plaque load and DAPI for untreated (A) and IBU-treated TG rats (B). Scale bar = 1000µm for panels (A) and (B). Plaque load was not reduced across hippocampal subregions collapsed (C), or in CA1, CA3 or SB (D,E, G). IBU-treatment significantly reduced plaque load in DG (**p < 0.01), (F).

### IBU significantly reduces tangle levels in the dentate gyrus

PHF1 staining was used to analyze tangle levels in the hippocampus of Tg-AD rats (Figs 3A-B). The results show that the levels of tangles were not significantly altered following treatment with IBU when collapsed across hippocampus subregions (Fig 3C; t = 0.4079, p = 0.6870). Analysis of individual subregions shows no significant difference between treatments in CA1 (Fig 3D t = 1.419, p = 0.1688), CA3 (Fig 3E; t = 0.9579, p = 0.3485), and SB (Fig 3G; t = 0.2824, p = 0.7807). Significant reduction in tangle levels was identified in dentate gyrus (DG) (Fig 3F; t = 7.295, p < 0.0001).

**Figure 3.**
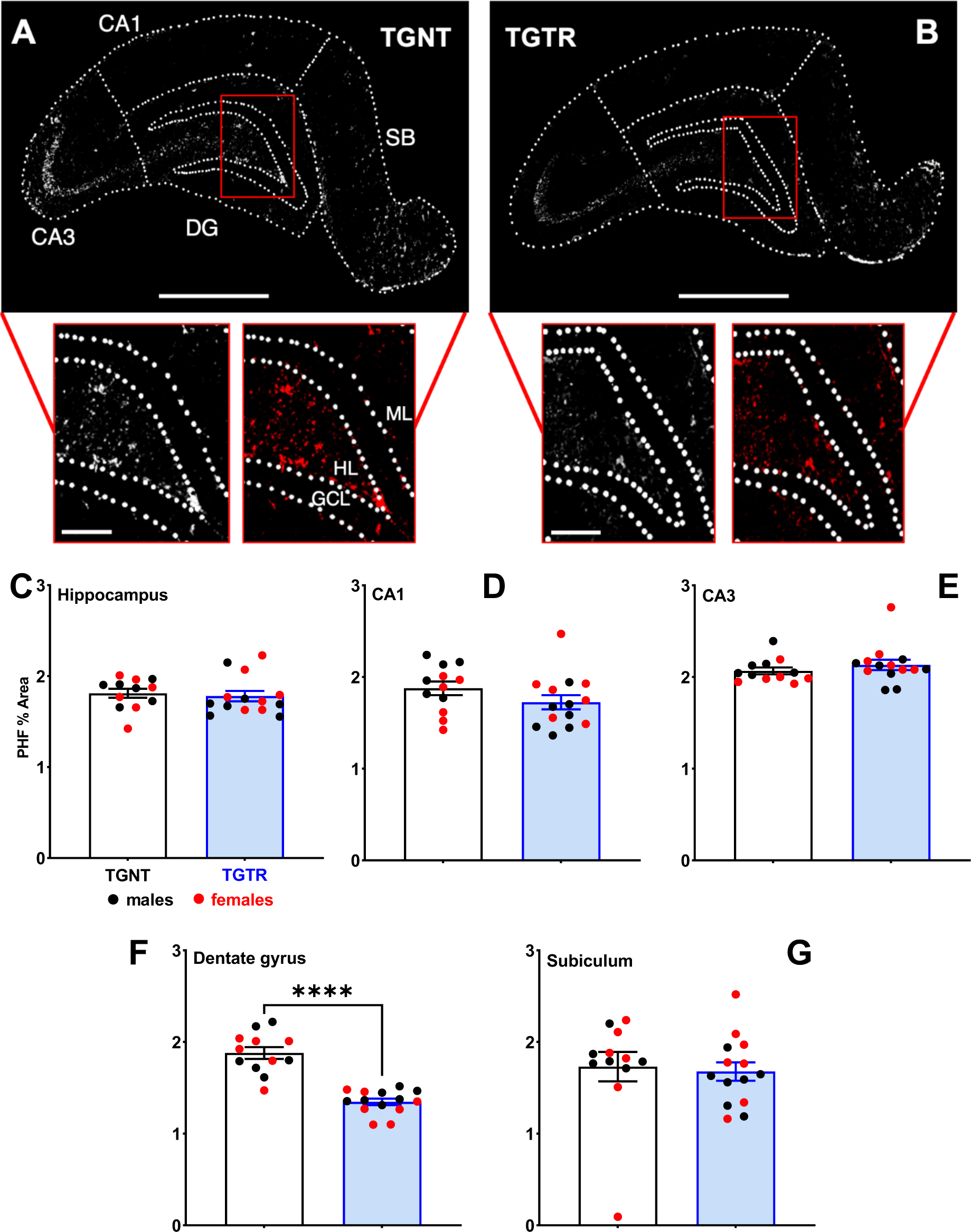
Analysis of tau paired helical filaments (PHF1) in Tg-AD rats. IHC for PHF1 staining across hippocampal subfields for TGNT (A) and TGTR rats (B). Scale bar = 1000 µm for (A) and (B) large panels, 250 µm for (A) and (B) small panels. PHF1 positive signal was not reduced across hippocampal subregions collapsed (C), in CA1 (D), CA3 (E), or Subiculum (G). IBU-treatment significantly reduced PHF1 signal in DG (****p < 0.0001) (F).

### IBU did not attenuate neuronal loss

We next examined whether IBU would affect the presence of healthy neurons through NeuN staining. WTNT (Fig 4A) and TGNT (Fig 4B) hippocampi show that TGNT has less NeuN staining compared to WTNT in the DG area (mean ± s.e.m.: 30.18 ± 0.10 % Area for WTNT, 25.74 ± 0.28 % Area; t = 15.10, p < 0.0001). The question to address was whether IBU-treatment could significantly reduce the degree of neuronal loss observed in TGNT. As with our prior staining, we analyzed the entire hippocampus collapsed across subregions, and then centered our focus on individual subregions. Our results showed that IBU-treatment did not significantly affect the degree of neuronal loss across the entire hippocampus (Fig 4C; t = 0.2237, p = 0.8250), nor within individual subregions CA1 (Fig 4D; t = 1.216, p = 0.2358), CA3 (Fig 4E; t = 1.498, p = 0.1474), DG (Fig 4F; t = 1.186, p = 0.2491), or SB (Fig 4G; t = 1.088, p = 0.2874).

**Figure 4.**
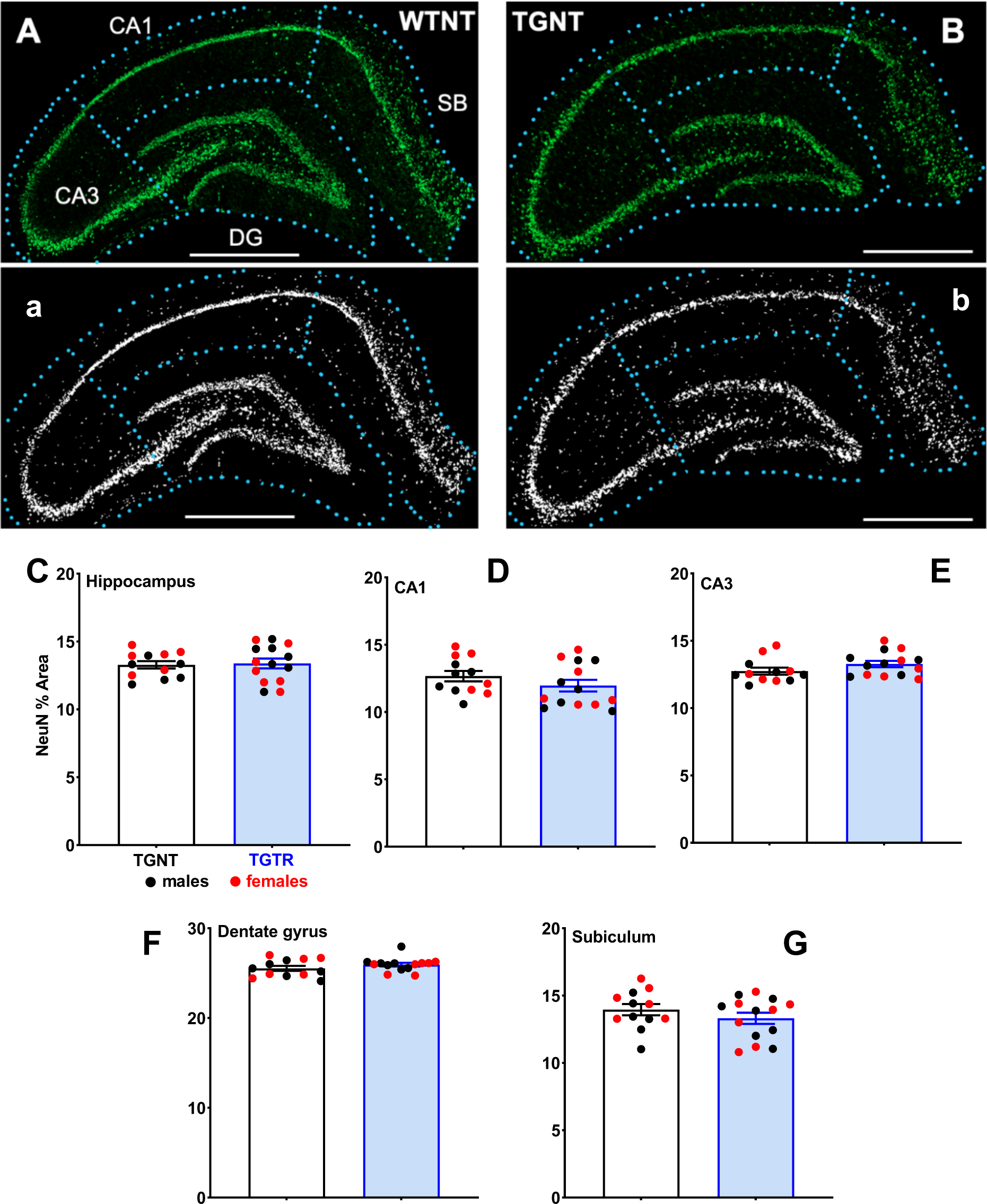
Neuronal loss across hippocampal subregions assessed with NeuN staining. DG shows a significant reduction in neuronal loss in TGNT vs WTNT, p < 0.0001 (A, B), corresponding masks shown in (a, b). There were no significant effects between TGNT vs TGTR with IBU on NeuN staining, neither collapsed across all subregions (C), nor in CA1 (D), CA3 (E), DG (F), or SB (G). Scale bar = 1000µm.

### IBU significantly reduces the ratio of amoeboid to ramified microglia in the hippocampus

Analysis for microglia using Iba1 staining shows that there are different forms that microglia can exhibit in the AD pathology. This pathology results in varying degrees of circularity of the microglia that is associated with a particular microglial type ranging from reactive to amoeboid (Fig 5A). Using this scale of circularity, we analyzed the ratio of amoeboid/ramified in the hippocampus and in the individual subregions. Our results show that in the hippocampus collapsed across subregions, there is a significant reduction in amoeboid/ramified ratio following IBU-treatment (F_(1,51)_ = 13.83, p < 0.01; Fig 5B) and significant post-hoc analyses between WTNT vs TGNT (t = 4.236; p < 0.01) and between TGNT and TGTR (t = 2.746; p = 0.0491). A similar pattern was shown in CA1 (F_(1,51)_ = 4.512, p = 0.0386; Fig 5C) and significant post-hoc analyses between WTNT vs TGNT (t = 4.044; p < 0.01) and between TGNT and TGTR (t = 2.946; p = 0.0289); and CA3 (F_(1,51)_ = 6.298, p = 0.0154; Fig 5D) and significant post-hoc analyses between WTNT vs TGNT (t = 3.114; p< 0.0182) and between TGNT and TGTR (t = 2.863; p = 0.0362). The DG subregion showed an overall effect of treatment (F _(1,51)_ = 16.32, p < 0.01; Fig 5E) and significant post-hoc analyses between WTNT vs TGNT (t = 3.931; p < 0.01). A similar pattern seen in DG was reflected in SB (F _(1,51)_ = 4.167, p = 0.0464; Fig 5F) and significant post-hoc analyses between WTNT vs TGNT (t = 2.779; p = 0.0448).

**Figure 5.**
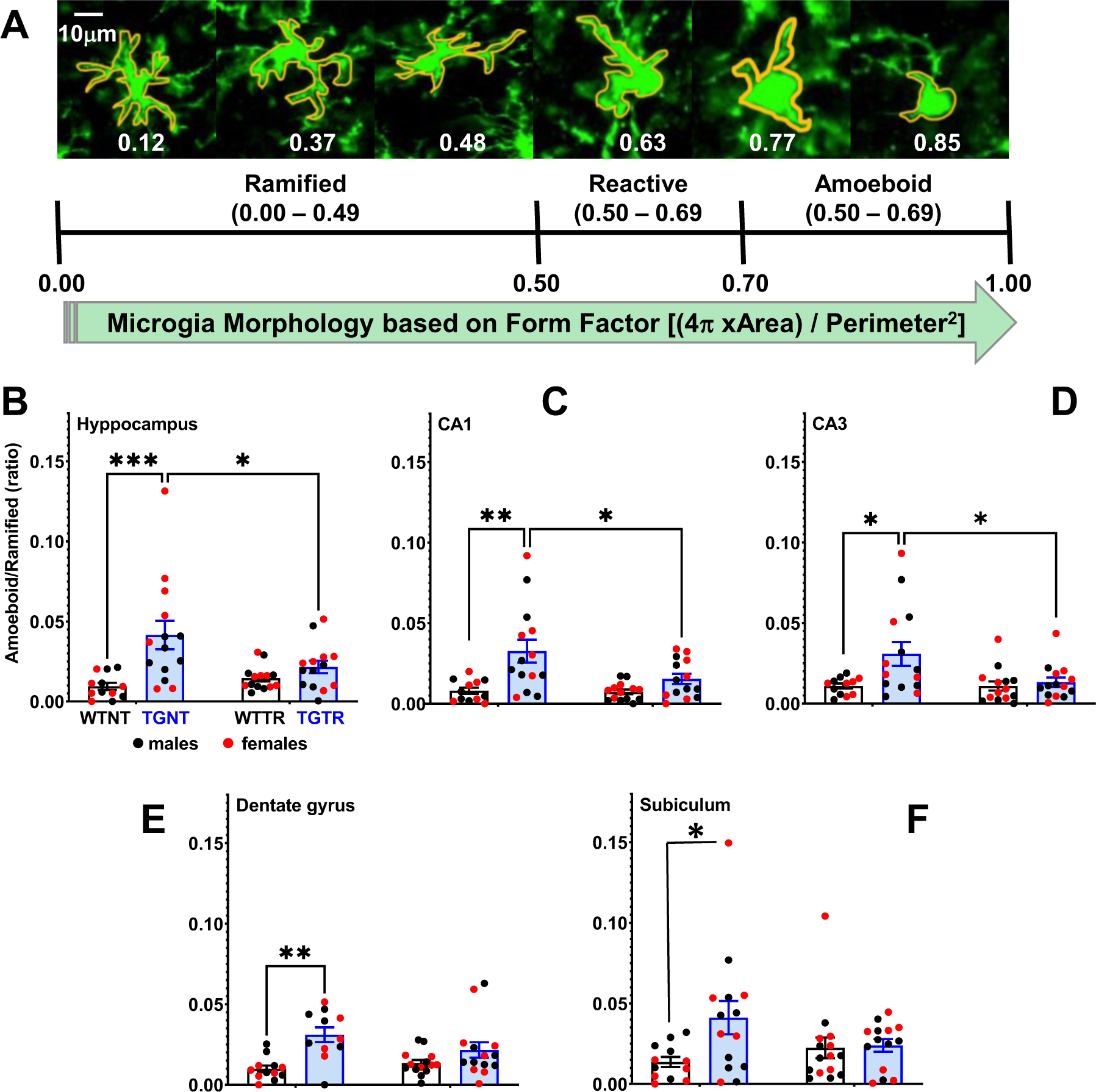
Microglial analyses following IBU-treatment. IHC for Iba1 staining identifying various form factors for circularity (A). Ratio for amoeboid/ramified expression collapsed across hippocampal subregions show significant post-hoc differences between WTNT vs TGNT and for individual subregions (CA1, CA3, Dentate gyrus and Subiculum (C-F). Significant post-hoc differences identified for TGNT vs TGTR in hippocampus collapsed across subregions, CA1, and CA3 (B-D). *p < 0.05; ** p < 0.01, ***p < 0.001.

### IBU affects gene expression levels of the TLR and ubiquitin-proteasome (UP) pathways differentially in males and females

Our hippocampal RNA sequencing analysis reported output measures as reads per million (RPM) for over 17,000 genes. Of particular interest are the effects IBU has on the mRNA expression of genes involved in TLR signaling, since IBU is a TLR4 antagonist. Thus, we compared hippocampal gene expression in the TGTR vs TGNT, separately in males and females (n = 5 per condition). Genes of interest included members of the IRAK family, such as IRAK isoforms, downstream targets of IRAK activation, and specific ubiquitin ligases and substrates (Fig. 6 and Supplemental Table S6), which eventually converge on the NF*κ*B transcription factor. Among the 16 genes of interest, five of them were unchanged in both sexes, including the adaptor MyD88, the ubiquitin (Ub) conjugase Ube2v1, and kinases IRAK 1, 2 and 4. The remaining 11 genes were either up or downregulated across sexes, or differentially expressed between them. The false discovery rate (FDR) and p-values are reported for the mRNA expression of the genes in terms of RPMs (Supplemental Table S6), and the gene expression for TGTR is shown as percent relative to TGNT (Figure 6).

**Figure 6.**
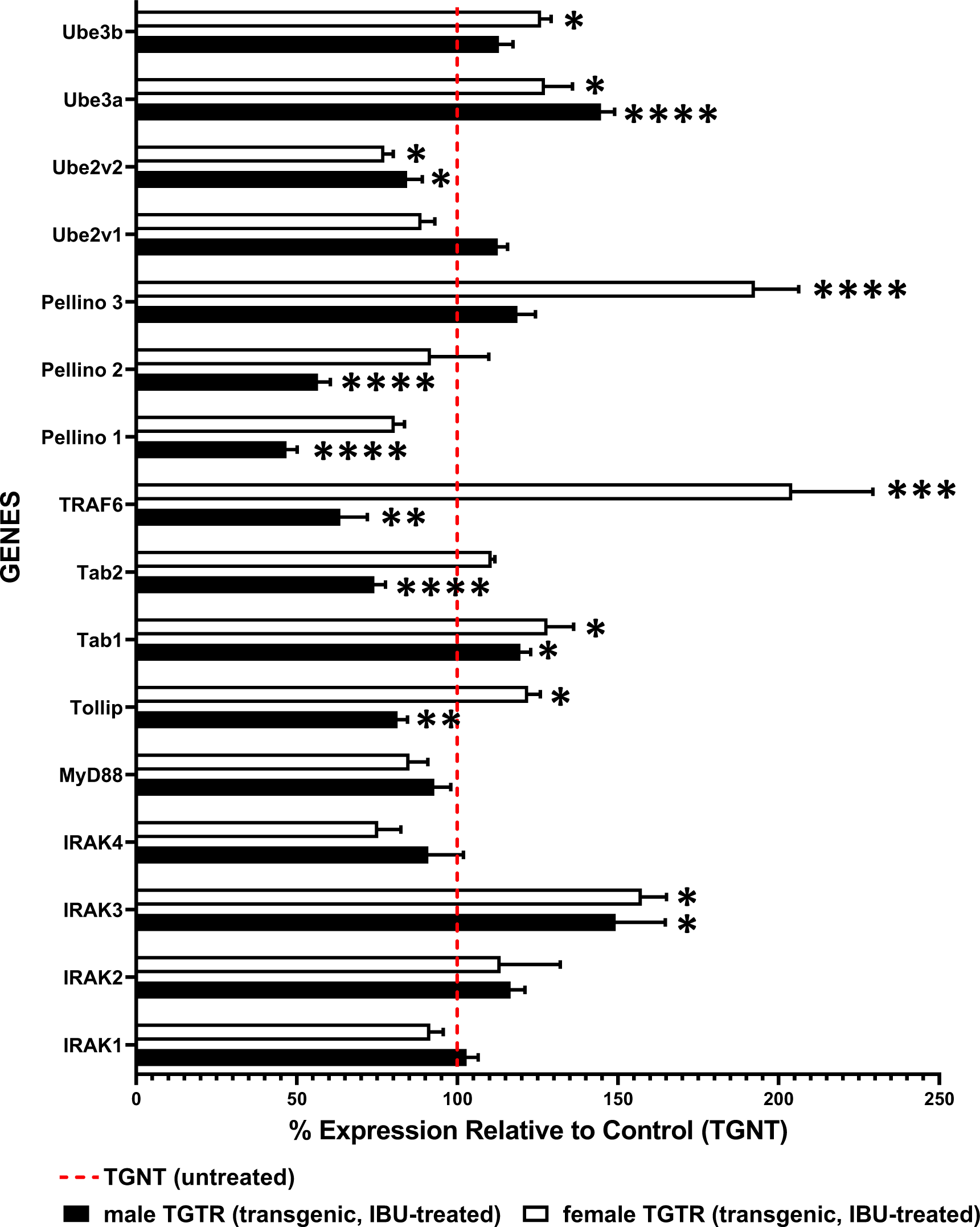
RNA sequencing data for TGTR and TGNT rats show differential expression of genes involved in the toll-like receptor (TLR) and Ub-proteasome (UP) pathways. These genes encode for IRAK-specific substrates, IRAK isoforms, and ubiquitin ligases and conjugating enzymes. Male and female hippocampal tissues were analyzed separately for mRNA expression in TGTR relative to TGNT. The dotted red line represents 100% of the gene expression for male or female TGNT rats. *p < 0.05; **p < 0.01, ***p < 0.001, ****p < 0.0001.

Upon IBU-treatment, we observed the following changes in TLR pathway gene expression: (1) IRAK3, a negative regulator of IRAK1, increased 1.50-fold in males and 1.57-fold in females. (2) IRAK1’s downstream target, the ubiquitin ligase Tumor-Necrosis Factor-Receptor Associated Factor 6 (TRAF6), decreased 1.57-fold in males, and increased 2.05-fold in females. (3) Similarly, Tollip, an IRAK1 substrate, decreased 1.23-fold in males, and increased 1.22-fold in females. (4) Other IRAK1 downstream targets such as TGF*β* Activated Kinase Binding Proteins (Tab 1 and 2), also showed significant changes. Tab1 increased 1.20-fold in males, and 1.28-fold in females, while Tab2 decreased 1.35 fold in males only.

Changes in the UP pathway gene expression were also detected upon IBU-treatment: (1) Pellino isoforms 1 and 2, which function as ubiquitin ligases, decreased in males: 2.14-fold for Pellino 1 and 1.76-fold for Pellino 2. Pellino 3, also a ubiquitin ligase, was upregulated in females by 1.93-fold. (2) Other ubiquitin ligases, such as Ube3a increased 1.45-fold in males and 1.27-fold in females, while Ube3b only increased in females (1.26-fold). (3) The ubiquitin conjugase variant 2 (Ube2v2) decreased 1.19-fold in males, 1.29-fold in females.

These findings show that IBU-treatment affects gene expression levels of the TLR and UP pathways differentially in males and females. Based on the TLR4 signaling pathway, we surmise that IBU-treatment inhibits IRAK1 activity by increasing expression of its negative regulator IRAK3, and/or by altering TRAF6 and other ubiquitin ligase and conjugase levels.

## Discussion

Due to the multifactorial nature of AD, conventional one-drug-one-target approaches are unfruitful in discovering effective AD therapeutics. The low success rate of target-based screening has revived interests in phenotypic screening based on cell-lines or animal models, since it could identify drug leads in a physiologically relevant condition. However, the phenotypic screening is relatively low-throughput, expensive, and difficult in the target deconvolution.

Furthermore, the drug response in the cell line or animal model can be significantly different from that in a patient’s tissue. The lack of mechanistic understanding of drug actions and a knowledge gap between a model system and an individual patient, makes it challenging to optimize drug lead compounds and to realize personalized medicine. In this study, we developed a multi-scale modeling approach that bridged target-based and phenotype-based drug repurposing. Our approach can not only use a patient’s tissue sample directly for the phenotype-based compound screening, but also deconvolute genome-wide drug-target interactions. We successfully identified IBU as a promising polypharmacological agent for the treatment of AD, demonstrating the potential for integrating machine learning, biophysics, and systems biology for the multi-scale modeling of drug actions. With the rapid advances in machine learning, especially deep learning techniques, we can further boost the performance of the multi-scale modeling and achieve personalized drug discovery.

We demonstrate for the first time in a transgenic rat model of AD (Tg-AD) that IBU-treatment mitigates cognitive deficits and hippocampal pathology associated with AD. This transgenic rat model developed by the Cohen and Town group^37^ is unique as it exhibits not only hippocampal-dependent spatial learning and memory deficits, but also hippocampal AD pathology including plaques, tau paired-helical filaments, neuronal loss and microgliosis, in a progressive age-dependent manner that mimics the pathology observed in AD patients. We chose to initiate IBU-treatment at an early age (5 months of age) based on previous studies with MS patients showing that the drug was effective only when administered in the early stage of the disease^22^. In our studies, the long-term daily IBU-treatment (6 months) was well tolerated by the Tg-AD rats as we did not identify any adverse effects. The IBU-treatment considerably improved spatial learning and memory and reduced AD pathology, although some of the AD-pathology was still detectable albeit at significantly lower levels. Overexpression of the human APPsw (2.6-fold) and PS1*Δ*E9 (6.2-fold) driving the robust pathology, may account for the incomplete protection provided by IBU. In addition, since AD is a multifactorial disorder, a combinatorial therapeutic approach is most likely required. The preventive effects of IBU-treatment that we observed in the Tg-AD rats, were most prominent in the DG region of the hippocampus, as the CA1 and CA3 regions showed less improvement. The reason for this hippocampal regional effect of IBU remains to be determined. However, since the DG is known to be vulnerable to aging and to be affected in the early stages of AD it is possible that the DG is also the most responsive to treatment^29 30^ . Overall, our data clearly demonstrate the potential of IBU to lessen deficits in spatial learning and memory, to mitigate plaque and tangle pathology, and to diminish neuronal loss and gliosis in the hippocampus of the Tg-AD rats.

A previous study supported the preventive effects of IBU-treatment in mice that received bilateral intracerebroventricular injections of Aβ1–42 ^31^. Pre-treatment for 15 days with IBU (i.p.) significantly ameliorated impaired spatial learning and memory, and inhibited hippocampal neuroinflammatory and apoptotic responses in the A*β*-injected mice. In an MPTP mouse model of Parkinson’s disease (PD), IBU was administered subcutaneously b.i.d. for 9 days, starting two days prior to 1-methyl-4-phenyl-1,2,3,6-tetrahydropyridine (MPTP) intoxication^32^. IBU diminished astroglial activation in the MPTP-treated mice, but had no impact on striatal dopaminergic cell survival seven days after acute MPTP intoxication. These two mouse studies evaluated the efficacy of a short-term IBU-treatment, in contrast to our studies that evaluated the effects of a long-term IBU-treatment using a transgenic rat model of AD that exhibits progressive age-dependent pathology. This is important because AD is a progressive neurodegenerative disorder with aging being the greatest risk factor. Thus, it is critical to evaluate the long-term effects of potential therapeutics.

IBU was first described as a non-specific PDE inhibitor, but more recently was shown to also act as a TLR4 antagonist^14 15^. IBU crosses the blood brain barrier, has a half-life of about 19h, improves blood flow to the brain, and protects against neuroinflammation^33^. The anti-inflammatory effects of IBU could be mediated by its PDE-inhibiting properties or by its TLR4 inactivation, and it is unclear which property potentiates the other, or if both work in synchrony. IBU-treatment exhibited immunomodulatory effects in MS and ALS by shifting the pro-inflammatory phenotype of microglia to an anti-inflammatory phenotype^34 35^. Based on these findings, we propose that the beneficial effects of IBU observed in our studies can be attributed to the anti-inflammatory effects of IBU.

The RNAseq analysis we conducted to compare gene expression in the hippocampus of untreated and IBU-treated transgenic rats, revealed that the TLR and UP pathways are altered by IBU-treatment. From the 16 genes we compared within the two pathways, five of them were not changed, including the adaptor MyD88, the Ub conjugase Ube2v1, and kinases IRAK 1, 2 and 4. The remaining 11 genes were either upregulated or downregulated. We will discuss two potential mechanisms that could mediate the anti-inflammatory action of IBU on the TLR pathway. For example, IBU-induced up-regulation of IRAK3, a central negative regulator of TLR signaling, prevents IRAK1 from activating TRAF6 through phosphorylation, thus preventing NF*κ*B activation. Furthermore, the TLR pathway is dependent on the function of the UP pathway at several steps, such as (1) degradation of IRAK1 after phosphorylation of TRAF6, (2) degradation of TRAF6 after activation of the TAK/TAB complex, and (3) degradation of I*κ*B, which is required for NF*κ*B nuclear translocation and activation. IBU-induced alterations on the expression of genes coding for components of the UPP, as shown in our data, indicate that this degradation pathway is recruited for modulation of inflammation via the TLR pathway.

The outcome and details of the IBU-induced alterations in the TLR and UP pathways will be addressed in future studies. However, their significance is crucial as IBU is a TLR4 antagonist.

Interestingly, TLR4-mediated induction of apoptosis is observed mostly in aging and not young neurons^36^. TLR4 mediated signaling, via still unclear mechanisms, seems to contribute to the pathology of age-related neurodegenerative diseases, including AD^36^. Using TLR4 antagonists, such as IBU, could offer an efficient means to prevent the damaging events associated with neuroinflammation in AD.

In Japan and other Asian countries, IBU is approved for treating asthma and stroke^33^. Our results suggest that IBU can be repurposed for use as a potential therapeutic to treat memory deficits and pathology in AD.

## Methods

### TgF-344AD transgenic rat model of AD and IBU treatment

Fisher transgenic 344-AD (Tg-AD) rats express mutant human *Swedish* amyloid precursor protein (APPsw) and *Δ* exon 9 presenelin-1 (PS1*Δ*E9) at 2.6-and 6.2-fold higher levels than the endogenous rat proteins. Expression of the two human mutant transgenes is driven by the prion promoter. No pathology differences were yet reported between sexes in this rat model of AD^37^.

The Tg-AD rats (males = 16; females = 13) and wild type (WT) (males = 15; females = 15) littermates were purchased from the Rat Resource and Research Center (RRRC, Columbia, MO), and arrived at our animal facility at Hunter College when they were approximately 4 weeks of age. The rats were housed in pairs on a 12h light/dark cycle with food and water available *ad libitum*. All animal procedures were approved by the Institutional Animal Care and Use Committee at Hunter College.

To investigate the therapeutic potential of IBU in Tg-AD rats, we opted to start drug treatment early at 5 months of age and continued treatment daily for 6 months, up to 11 months of age, when the Tg-AD rats exhibit most of the full AD-like pathology. IBU (cat # HY-B0763, MCE, Monmouth Junction, NJ) was administered orally combined with rodent chow obtained from Research Diets Inc. (NJ), 10 mg / kg body weight (Supplemental Figures S1 and S2). Rats were analyzed for cognitive behavior at 11 months of age, prior to sacrificing. Rat brains were analyzed for immunohistochemical (IHC) and RNAseq analyses.

### Cognitive behavior assessment

*Active Place Avoidance Task (aPAT)* is a spatial learning and memory test that assesses hippocampal function^38^. The arena is comprised of a rotating platform that operates at one revolution per minute (rpm), with visual cues at the four walls of the room. The arena is divided into four quadrants, one of which will be assigned as a shock zone. If a rat enters the shock zone and remains there for at least 1.5 seconds, it will receive a 0.2 amp shock. A shock is given every 1.5 seconds until the rat leaves the shock zone. An overhead camera is perched atop of the arena to track the location of the rat as it performs the task.

The rats were given a 30-min resting period in a paper-bedding cage prior to the habituation trial. During the 10-min *<u>habituation trial</u>*, rats were permitted to explore while the shock zone is off. Afterwards, the shock zone was turned on and rats begun their training, which consisted of six 10-min *<u>training trails</u>*, with a 10-min inter trial interval in the home cage. 24 hours after the last training trial, rats were given one 5-min *<u>test trial</u>* with the shock zone turned off.

### Immunohistochemical analysis (IHC) for Aβ Plaques and Microglia

At 11 months of age, the rats were anesthetized (i.p.) with ketamine (100 mg/kg) and xylazine (10 mg/kg), and transcardially perfused for 15-min with cold RNAse-free 1X-PBS. The rat brains were removed and the right hemisphere was micro-dissected (regions: prefrontal cortex, cingulate cortex, entorhinal cortex, and hippocampus) and immediately snap frozen for later RNAseq analysis. The left hemispheres were sequentially post-fixed with 4% paraformaldehyde for 48 hours at 4°C, cryoprotected in a 30% sucrose/PBS solution at 4°C until they sank to the bottom of the vial, flash frozen in 2-methybutane, and stored at -80°C until sectioned. The left hemispheres were sectioned with a Leica CM 3050S cryostat. Coronal sections, 30µm in thickness, were collected serially along the anteroposterior axis and stored at -20°C in cryoprotectant (30% glycerol and ethylene glycol in 1X PBS) until use.

Hippocampal sections for these IHC studies were located between -3.36 mm to -4.36 mm relative to bregma. Sections were processed with a mounted protocol for IHC analyses as described previously^39^. After mounting, hippocampal sections were immersed in 0.05 M glycine (Fisher BioReagents cat# BP3815) in 0.3% TritonX (Thermo Fisher, cat# PI85112) in 1X PBS (T-PBS) for 30-min to reduce autofluorescence, and then sequentially rinsed in 0.3% Triton-PBS, incubated in 15% normal goat serum (NGS) in 0.3% Triton-PBS blocker for 30-min, and incubated with primary antibody cocktail containing 15% NGS and 0.03% T-PBS overnight at 4°C on a rocker. Primary antibodies were diluted at 1:1000 for A*β* 4G8 antibody (epitope a.a.

17-24) (Biolegend – mouse – cat# 800708) and 1:500 for microglia Iba1 (Wako – rabbit – cat# 019-19741). The following day, sections were washed in 0.03% Triton-PBS, and incubated in fluorescent secondary antibody cocktail 15% NGS and 0.03% Triton-PBS for 2 hours.

Secondary antibodies included Alexa Fluor goat anti-mouse 568 (cat# A-11031, Life Technologies, Thermo Fisher Sci.) and Alexa Fluor goat anti-rabbit 488 (cat# A-11008, Life T-PBS, then PBS, and finally cover-slipped with VectaShield^®^ mounting media with DAPI (#sku H-1200-10). Slides were kept at 4°C in the dark until imaged.

To establish the significance of the effects of IBU on AD-related pathology, the IHC images were quantified. Hippocampal subfields (CA1, CA3, DG, and SB) were isolated, cropped, and saved as .tif files for use in pixel-intensity area analyses as described previously, along with analyzing the overall hippocampus^39^. Images were analyzed to extract the positive signal from each image with custom batch-processing macroscripts created for each channel/marker. Pixel-intensity statistics were calculated from the images at 16-bit intensity bins. Positive signal within each cropped image were extracted using the following formulae: average pixel intensity + [(1.25 *[Iba1]* or 2.0 *[Aβ]*) x Standard deviation of intensities]^40^. Positive signal were then measured, masks created, and merged when co-localization analyses was required.

### Microglia Analysis

Activated microglia exhibit a variety of morphologies that can be associated with their functions, distributed into three different groups according to their form factor (FF) (Figure 5A) which is defined as 4π X area/perimeter^2 41^. Each of the three microglia groups is defined as follows: *Ramified*, FF: 0 to 0.49; which actively engage in neuronal maintenance providing neurotrophic factors. *Reactive*, FF: 0.50 to 0.69; which are responsive to CNS injury, and *amoeboid*, FF 0.70 to 1; amorphous with pseudopodia. Microglia within each cropped Iba1 image were extracted using the following formula: average pixel intensity + [1.5 x standard deviation of intensities], and particles within 50–800 µm^2^ were chosen for FF analyses. Quantities of each microglia class were collected and analyzed in ratios to each other, and then compared across treatment/genotype. For this analysis, three tissue sections per treatment/genotype group were used for quantification. Nonspecific background density was corrected using ImageJ rolling-ball method^41^. Sections chosen for analysis corresponded to figures 60 – 70 of the rat brain atlas (bregma - 3.36 mm through - 4.44 mm)^42^.

### Immunohistochemical Analysis for NeuN and PHF1

Immunohistochemical analysis for NeuN and PHF1 was conducted following a similar protocol for A*β* and Iba1, with the following changes: 0.3% TritonX (Thermo Fisher, cat# PI85112) in 1X TBS (TBS-T) was used for all wash steps. Dry blocking milk (5% by weight) in 0.3% TBS-T was the blocking buffer^43^. The primary antibody cocktail was also prepared using the blocking buffer. Primary antibodies were diluted at 1:250 for PHF1 antibody (courtesy of the late Dr. Peter Davies) and 1:250 for NeuN (Millipore – chicken – cat# ABN91). The secondary antibody cocktail was composed of 20% Superblock/TBS in 0.03% TBS-T. Secondary antibodies included Alexa Fluor IgG1 goat anti-mouse 568 (1:80 dilution, cat# A-21124, Life Technologies, Thermo Fisher Sci.) and Alexa Fluor goat anti-chicken 488 (1:250 dilution, cat# 11039, Life Technologies, Thermo Fisher Sci.). Sections were washed with 0.03% TBS-T and TBS. Slides were kept at 4°C in the dark until imaged.

### PHF1 and NeuN Analysis

Hippocampal subfields (CA1, CA3, DG, and SB) were isolated, cropped, and saved as .tif files for use in pixel-intensity area analyses as described previously, along with analyzing the overall hippocampus^39^. Images were analyzed to extract the positive signal from each image with custom batch-processing macroscripts created for each channel/marker. Pixel-intensity statistics were calculated from the images at 16-bit intensity bins. Positive signal within each cropped image was extracted using the following formulae: average pixel intensity + [(2.0 [*PHF1]* or 1.25 *[NeuN]*) x Standard deviation of intensities]^40^. Positive signals were then measured, masks created, and merged when co-localization analyses was required.

### RNAseq analysis

RNAseq analyses were performed using the outsourcing service at the UCLA Technology Center for Genomics & Bioinformatics. Briefly, total RNA was isolated from the hippocampi of IBU-treated and untreated transgenic rats using the RNeasy Mini Kit (Qiagen cat# 74104). The integrity of total RNA was examined by the Agilent 4200 TapeStation System. Libraries for RNAseq were constructed with the Kapa Stranded mRNA Kit (Roche, cat. # KK8421) to generate strand-specific RNAseq libraries. Libraries were amplified and sequencing was performed with the HiSeq3000 sequencer. Gene expression data were normalized as the reads count per million (RPM) using the TMM method. Differentially expressed genes from IBU-treated transgenic rats were determined using the edgeR program^44^. The data generated are shown as the fold-change, p-value, and FDR for each gene.

### Kinase Binding Assay

To validate our computational predictions, we employed a competition biding assay to detect the binding of IBU to 425 human kinases. The proprietary KINOMEScan^TM^ was performed by Eurofins/DIscoverX (Fremont, CA). The tests were performed at 10 µM and 100 µM concentrations of levosimendan, respectively. Assay results were reported as % Control, calculated as follows:

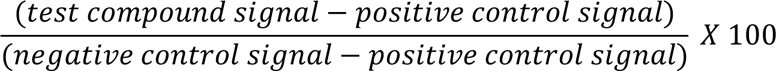

A lower % Control score indicates a stronger interaction. The KINOMEScanTM experiment and data analysis were performed by Eurofins/DiscoverX (Fremont, CA).

### Multi-scale predictive modeling of drug action

-Derivation of AD Signature:

We used six microglial RNAseq. Samples were obtained from ROSMAP projects through AMP-AD Knowledge Portal^45^. There were four samples from AD patients and two normal controls.

The sample IDs are shown in supplemental Table S5. We used Salmon^46^ quantify, the transcripts and DEseq2^47^ to perform differential expression analysis. Top differential expressed genes with adjust p value < 0.05 are used as AD signatures for further analysis.

-Phenotypic compound screening:

The up and downregulated genes in AD signature who also served as LINCS L1000 landmark genes were used to screen drug-induced expression profiles in L1000 dataset. The drug-induced expression profiles we used here are processed by Bayesian signature detection pipeline^48^. They are compared to the AD signature by Gene Set Enrichment Analysis (GSEA)^49^, and drugs with the lowest scores are selected as candidates.

-Prediction of genome-wide drug-target interactions:

We applied 3D-REMAP to predict drug off-target binding of IBU. The details of 3D-REMAP were published elsewhere. In brief, 3D-REMAP first represents annotated chemical-protein interactions (CPIs) between millions of chemicals and thousands of proteins into a matrix. Then it applies structure-bases ligand binding similarity searches and protein-ligand docking to fill missing entries in the CPI matrix. Finally, it uses a one-class collaborative filtering algorithm to predict all unknown CPIs.

The details of methods were published elsewhere^28^. In brief, disease, gene, and chemical name entries were recognized using DNorm v0.0.6^50^, Gimli v1.0.2^51^, and tmChem^52^, respectively.

Word2Vec^27^ was used to map each processed term in the PubMed abstracts to a 300-dimensional vector, which represents that term’s relation to other terms. Drug-gene-disease associations were then evaluated by the cosine similarity between the vector representation of drug, gene, and disease terms.

-Prediction of BBB permeant:

The BBB permeant of chemical compounds was predicted using SwissADME^53^.

## Acknowledgements

We thank the late Dr. Peter Davies (Albert Einstein University, NY, NY) for generously providing the PHF1 antibody. This work was supported in part by NIH/NIGMS R01GM122845 and NIH/NIA R01AG057555 to LX and NIGMS Training Grant RISE 5R25GM060665 to GO and OC, and the City University of New York (Ph.D. program in Biochemistry, Graduate Center).

## Author Contributions

QL, YQ, and LX performed computational modeling and identified Ibudilast as treatment. MFP, PR and PS designed experiment. LX designed algorithms and computational analysis. GO, PR, MFP, PS, and LX wrote the manuscript. GO and CW collected the weekly animal weights and performed the behavioral assays, and statistical analyses. GO and OC conducted the IHC and analyses. All authors approved the manuscript for submission.

## Competing Interest

The authors declare that the research was conducted in the absence of any commercial or financial relationships that could be construed as a potential conflict of interest.

## Data Availability

The raw data supporting the conclusions of this manuscript will be made available upon publication.

## Supplemental Tables and Figures

Table S1. Differential gene expression profile of AD patients’ microglia vs normal control.

Table S2. Top 100 drugs identified from L1000 drug signature analysis.

Table S3. IBU KinomeScan^TM^ results under 10 μM of IBU.

Table S4. IBU KinomeScan^TM^ results under 100 μM of IBU.

Table S5. ROSMAP samples used to derive AD disease signature.

Table S6. RNA sequence analysis for selected TLR and UP pathway genes.

Figure S1. IBU dosage during experimental period.

Figure S2. Weight change during the experimental period for WTTR and TGTR males and females.

## Notes

### Competing Interest Statement

The authors have declared no competing interest.

## References

1. 2020 Alzheimer’s disease facts and figures. Alzheimers Dement (2020).

2. Serrano-Pozo, A., Frosch, M.P., Masliah, E. & Hyman, B.T. Neuropathological alterations in Alzheimer disease. Cold Spring Harb Perspect Med 1, a006189 (2011).

3. Mufson, E.J., Malek-Ahmadi, M., Perez, S.E. & Chen, K. Braak staging, plaque pathology, and APOE status in elderly persons without cognitive impairment. Neurobiol Aging 37, 147–153 (2016).

4. Mehta, D., Jackson, R., Paul, G., Shi, J. & Sabbagh, M. Why do trials for Alzheimer’s disease drugs keep failing? A discontinued drug perspective for 2010-2015. Expert Opin Investig Drugs 26, 735–739 (2017).

5. Amor, S., Puentes, F., Baker, D. & van der Valk, P. Inflammation in neurodegenerative diseases. Immunology 129, 154–169 (2010).

6. Crisafulli, S.G., Brajkovic, S., Cipolat Mis, M.S., Parente, V. & Corti, S. Therapeutic Strategies Under Development Targeting Inflammatory Mechanisms in Amyotrophic Lateral Sclerosis. Mol Neurobiol 55, 2789–2813 (2018).

7. Barkhof, F., et al. Ibudilast in relapsing-remitting multiple sclerosis: a neuroprotectant? Neurology 74, 1033–1040 (2010).

8. Tiao, G., et al. Sepsis stimulates nonlysosomal, energy-dependent proteolysis and increases ubiquitin mRNA levels in rat skeletal muscle. J Clin Invest 94, 2255–2264 (1994).

9. Bruno, K., et al. Targeting toll-like receptor-4 (TLR4)-an emerging therapeutic target for persistent pain states. Pain 159, 1908–1915 (2018).

10. Wang, L., et al. Crystal structure of human IRAK1. Proc Natl Acad Sci U S A 114, 13507–13512 (2017).

11. Rhyasen, G.W. & Starczynowski, D.T. IRAK signalling in cancer. Br J Cancer 112, 232–237 (2015).

12. Kim, J.H., et al. Pellino-1, an adaptor protein of interleukin-1 receptor/toll-like receptor signaling, is sumoylated by Ubc9. Mol Cells 31, 85–89 (2011).

13. Hutchinson, M.R., et al. Reduction of opioid withdrawal and potentiation of acute opioid analgesia by systemic AV411 (ibudilast). Brain Behav Immun 23, 240–250 (2009).

14. Jia, Z.J., Wu, F.X., Huang, Q.H. & Liu, J.M. [Toll-like receptor 4: the potential therapeutic target for neuropathic pain]. Zhongguo Yi Xue Ke Xue Yuan Xue Bao 34, 168–173 (2012).

15. Ain, Q.U., Batool, M. & Choi, S. TLR4-Targeting Therapeutics: Structural Basis and Computer-Aided Drug Discovery Approaches. Molecules 25(2020).

16. Ohashi, M., Uno, T. & Nishino, K. Effect of ibudilast, a novel antiasthmatic agent, on anaphylactic bronchoconstriction: predominant involvement of endogenous slow reacting substance of anaphylaxis. Int Arch Allergy Immunol 101, 288–296 (1993).

17. Ruiz-Perez, D., et al. The Effects of the Toll-Like Receptor 4 Antagonist, Ibudilast, on Sevoflurane’s Minimum Alveolar Concentration and the Delayed Remifentanil-Induced Increase in the Minimum Alveolar Concentration in Rats. Anesth Analg 122, 1370–1376 (2016).

18. Lee, J.Y., et al. Ibudilast, a phosphodiesterase inhibitor with anti-inflammatory activity, protects against ischemic brain injury in rats. Brain Res 1431, 97–106 (2012).

19. Yamazaki, T., Anraku, T. & Matsuzawa, S. Ibudilast, a mixed PDE3/4 inhibitor, causes a selective and nitric oxide/cGMP-independent relaxation of the intracranial vertebrobasilar artery. Eur J Pharmacol 650, 605–611 (2011).

20. Suzumura, A., Ito, A., Yoshikawa, M. & Sawada, M. Ibudilast suppresses TNFalpha production by glial cells functioning mainly as type III phosphodiesterase inhibitor in the CNS. Brain Res 837, 203–212 (1999).

21. Fox, R.J., et al. Phase 2 Trial of Ibudilast in Progressive Multiple Sclerosis. N Engl J Med 379, 846–855 (2018).

22. Fox, R.J., et al. Design, rationale, and baseline characteristics of the randomized double-blind phase II clinical trial of ibudilast in progressive multiple sclerosis. Contemp Clin Trials 50, 166–177 (2016).

23. Pham, T.-H., Qiu, Y., Zeng, J., Xie, L. & Zhang, P. A deep learning framework for high-throughput mechanism-driven phenotype compound screening and its application to COVID-19 drug repurposing. Nature Machine Intelligence 3, 247–257 (2021).

24. Hodes, R.J. & Buckholtz, N. Accelerating Medicines Partnership: Alzheimer’s Disease (AMP-AD) Knowledge Portal Aids Alzheimer’s Drug Discovery through Open Data Sharing. Expert Opin Ther Targets 20, 389–391 (2016).

25. Lim, H., et al. Rational discovery of dual-indication multi-target PDE/Kinase inhibitor for precision anti-cancer therapy using structural systems pharmacology. PLoS Comput Biol 15, e1006619 (2019).

26. Karch, C.M., Cruchaga, C. & Goate, A.M. Alzheimer’s disease genetics: from the bench to the clinic. Neuron 83, 11–26 (2014).

27. Mikolov, T, Chen, K., Corrado, G., Dean, J. Efficient Estimation of Word Representations in Vector Space. arXiv:1301.3781v3. (2013).

28. Dider, S., Ji, J., Zhao, Z. & Xie, L. Molecular mechanisms involved in the side effects of fatty acid amide hydrolase inhibitors: a structural phenomics approach to proteome-wide cellular off-target deconvolution and disease association. NPJ Syst Biol Appl 2, 16023 (2016).

29. Noguchi, M., Mori, A., Sakamoto, K., Nakahara, T. & Ishii, K. Vasodilator effects of ibudilast on retinal blood vessels in anesthetized rats. Biol Pharm Bull 32, 1924–1927 (2009).

30. Takeda A, Tamano H. Is Vulnerability of the Dentate Gyrus to Aging and Amyloid-β1 42 Neurotoxicity Linked with Modified Extracellular Zn^2+^ Dynamics? Biol Pharm Bull. 2018;41(7):995–1000. doi: 10.1248/bpb.b17-00871. PMID: 29962410.

31. Wang, H., et al. Pretreatment with antiasthmatic drug ibudilast ameliorates Abeta 1-42-induced memory impairment and neurotoxicity in mice. Pharmacol Biochem Behav 124, 373–379 (2014).

32. Schwenkgrub, J., et al., The phosphodiesterase inhibitor, Ibudilast attenuates neuroinflammations in the MPTP model of Parkinson’s disease. PLoS ONE 12(7). e0182019.

33. Johnson, K.W., et al. Ibudilast for the treatment of drug addiction and other neurological conditions. Clin. Invest. (2014) 4(3), 269–279.

34. Khalid, S.I., Ampie, L., Kelly, R., Ladha, S.S. & Dardis, C. Immune Modulation in the Treatment of Amyotrophic Lateral Sclerosis: A Review of Clinical Trials. Front Neurol 8, 486 (2017).

35. Appel, S.H., Zhao, W., Beers, D.R., Henkel, J.S. The Microglial-Motoneuron dialogue in ALS. Acta Myologica, 4–8 (2011).

36. Calvo-Rodriguez, M., Garcia-Rodriguez, C., Villalobos, C. & Nunez, L. Role of Toll Like Receptor 4 in Alzheimer’s Disease. Front Immunol 11, 1588 (2020).

37. Cohen, R.M., et al. A transgenic Alzheimer rat with plaques, tau pathology, behavioral impairment, oligomeric abeta, and frank neuronal loss. J Neurosci 33, 6245–6256 (2013).

38. Lesburgueres, E., Sparks, F.T., O’Reilly, K.C. & Fenton, A.A. Active place avoidance is no more stressful than unreinforced exploration of a familiar environment. Hippocampus 26, 1481–1485 (2016).

39. Avila, J.A., et al. PACAP27 mitigates an age-dependent hippocampal vulnerability to PGJ2-induced spatial learning deficits and neuroinflammation in mice. Brain Behav 10, e01465 (2020).

40. Kerfoot, E.C., Agarwal, I., Lee, H.J. & Holland, P.C. Control of appetitive and aversive taste-reactivity responses by an auditory conditioned stimulus in a devaluation task: a FOS and behavioral analysis. Learn Mem 14, 581–589 (2007).

41. Dahlmann, B., Kuehn, L., Grziwa, A., Zwickl, P. & Baumeister, W. Biochemical properties of the proteasome from Thermoplasma acidophilum. Eur J Biochem 208, 789–797 (1992).

42. Nie, B., et al. A rat brain MRI template with digital stereotaxic atlas of fine anatomical delineations in paxinos space and its automated application in voxel-wise analysis. Hum Brain Mapp 34, 1306–1318 (2013).

43. Morrone, C.D., et al. Regional differences in Alzheimer’s disease pathology confound behavioural rescue after amyloid-beta attenuation. Brain 143, 359–373 (2020).

44. Robinson, M.D., McCarthy, D.J. & Smyth, G.K. edgeR: a Bioconductor package for differential expression analysis of digital gene expression data. Bioinformatics 26, 139–140 (2009).

45. De Jager, P.L., et al. A multi-omic atlas of the human frontal cortex for aging and Alzheimer’s disease research. Sci Data 5, 180142 (2018).

46. Patro, R., Duggal, G., Love, M.I., Irizarry, R.A. & Kingsford, C. Salmon provides fast and bias-aware quantification of transcript expression. Nat Methods 14, 417–419 (2017).

47. Love, M.I., Huber, W. & Anders, S. Moderated estimation of fold change and dispersion for RNA-seq data with DESeq2. Genome Biol 15, 550 (2014).

48. Qiu, Y., Lu, T., Lim, H. & Xie, L. A Bayesian approach to accurate and robust signature detection on LINCS L1000 data. Bioinformatics 36, 2787–2795 (2020).

49. Subramanian, A., et al. Gene set enrichment analysis: a knowledge-based approach for interpreting genome-wide expression profiles. Proc Natl Acad Sci U S A 102, 15545–15550 (2005).

50. Leaman, R., Islamaj Dogan, R. & Lu, Z. DNorm: disease name normalization with pairwise learning to rank. Bioinformatics 29, 2909–2917 (2013).

51. Campos, D., Matos, S., Oliveria, J.L., Gimil: open source and high performance biomedical name recognition. BMC Bioinformatics, 14–54 (2013).

52. Leaman, R., Wei, C.H., Lu, Z. tmChem: a high performance application for named entity recognition and normalization. Journal of Cheminformatics 7(Suppl 1):S3 (2015).

53. Daina, A., Michielin, O. & Zoete, V. SwissADME: a free web tool to evaluate pharmacokinetics, drug-likeness and medicinal chemistry friendliness of small molecules. Sci Rep 7, 42717 (2017).

